# Who is this gene and what does it do? A toolkit for munging transcriptomics data in python

**DOI:** 10.1101/299107

**Authors:** Charles K. Fisher, Aaron M. Smith, Jonathan R. Walsh

**Affiliations:** Unlearn.AI, Inc., San Francisco, CA 94108

## Abstract

Transcriptional regulation is extremely complicated. Unfortunately, so is working with transcriptional data. Genes can be referred to using a multitude of different identifiers and are assigned to an ever increasing number of categories. Gene expression data may be available in a variety of units (e.g, counts, RPKMs, TPMs). Batch effects dominate signal, but metadata may not be available. Most of the tools are written in R. Here, we introduce a library, genemunge, that makes it easier to work with transcriptional data in python. This includes translating between various types of gene names, accessing Gene Ontology (GO) information, obtaining expression levels of genes in healthy tissue, correcting for batch effects, and using prior knowledge to select sets of genes for further analysis. Code for genemunge is freely available on Github.

## I. OVERVIEW

munge: verb

1. to manipulate (raw data), especially to convert (data) from one format to another.

www.dictionary.com/browse/munge

Like any area that uses big data, transcriptomics data requires extensive munging – rote but critical tasks such as cleaning data, selecting relevant data, structuring metadata, and making labels interpretable. These tasks often need to be repeated on a given project as the data and aims evolve, and tend to be similar between different analyses. To face these challenges, a library of data munging tools can be extraordinarily useful. Such a library can provide reliable and tested tools to cleanly separate munging tasks from analysis, making it easier to start new projects and data processing pipelines less fragile.

This note introduces `genemunge`, a library of tools for working with human transcriptomics data. `genemunge` is written in python and is available as a package through `PyPI`. This initial version, `v0.0`, contains tools for tasks such as:

- Translating between conventions for gene symbols [1].
- Accessing Gene Ontology (GO) metadata [2–4].
- Using prior knowledge of biological and molecular processes to select gene sets [5, 6].
- Retrieving statistics on gene expression in healthy tissue [7–10].
- Converting expression data to TPM from counts or RPKM [11].
- Correcting for unobserved, and uninteresting, factors of variation (i.e., batch effects). [12, 13].

The goal of `genemunge`, and its current use case for the authors, is to serve as a resource for gene information that can return useful data structures and be integrated into processing pipelines. The next section gives a few example use cases of the library.

## II. EXAMPLE USE CASES OF GENEMUNGE

We consider a simple analysis where `genemunge` is useful. Suppose we want to find genes associated with the immune system, select those with larger expression in the small intestine than the stomach, and then retrieve basic information about those genes.

The following code snippets use the API from `genemunge v0.0`. We begin by importing libraries required for the example.

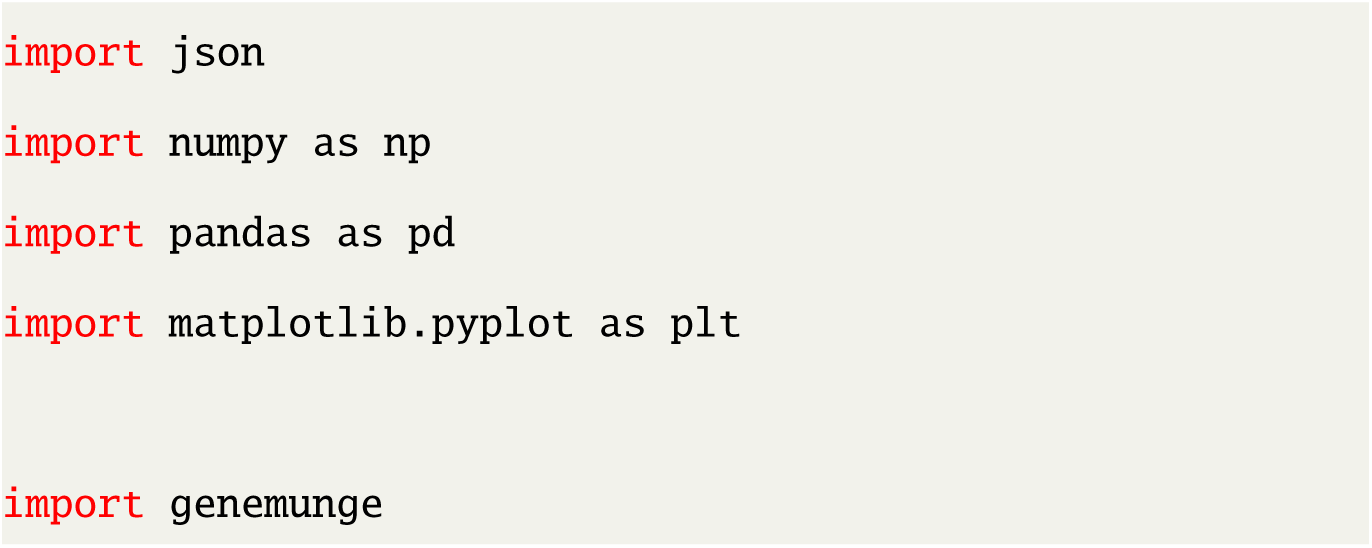

### A. Searching the gene ontology for relevant genes

The Gene Ontology (GO) contains basic descriptors for each ontology entry [2–4]. Since we want genes related to the immune system, we will do a keyword search for *immune* and retrieve GO identifiers with this keyword. We can then obtain the associated genes.

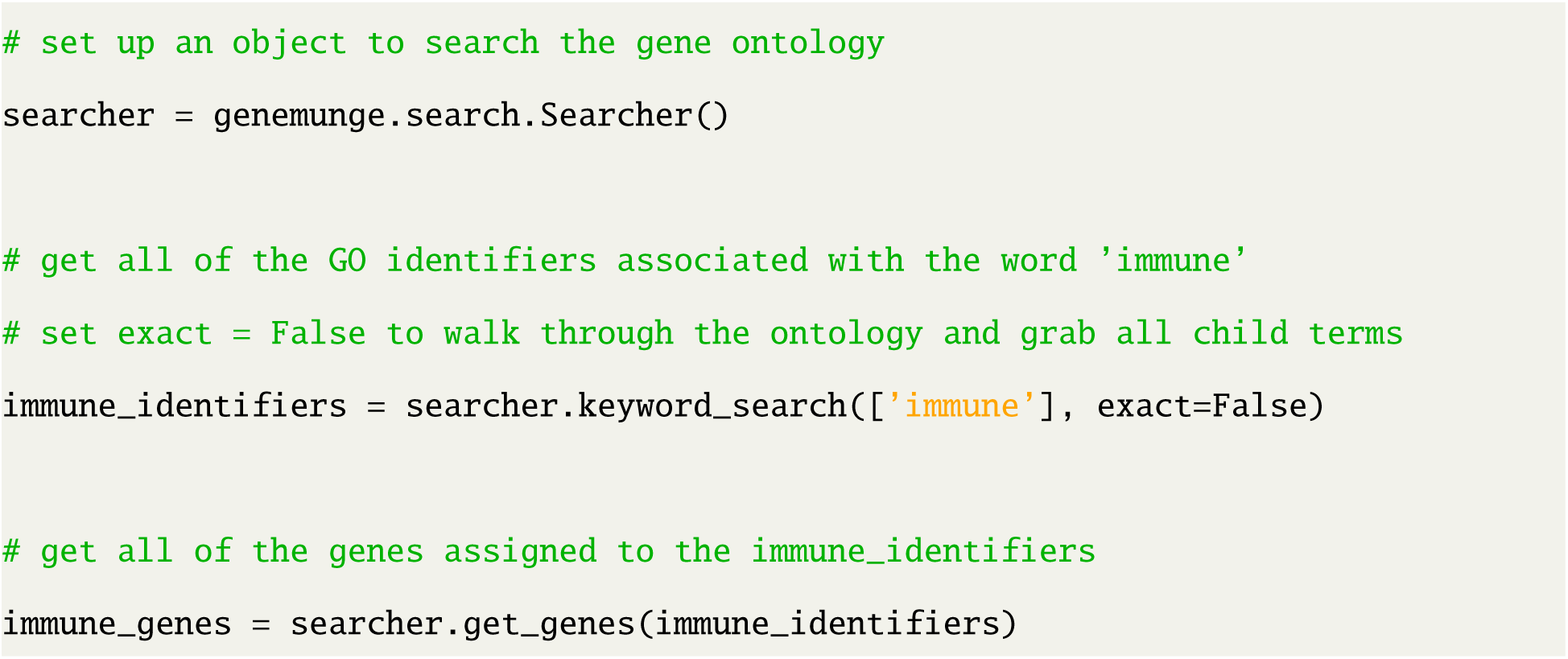

We will then use prior knowledge to remove housekeeping genes. A list of housekeeping genes curated by [6] is stored in `genemunge`.

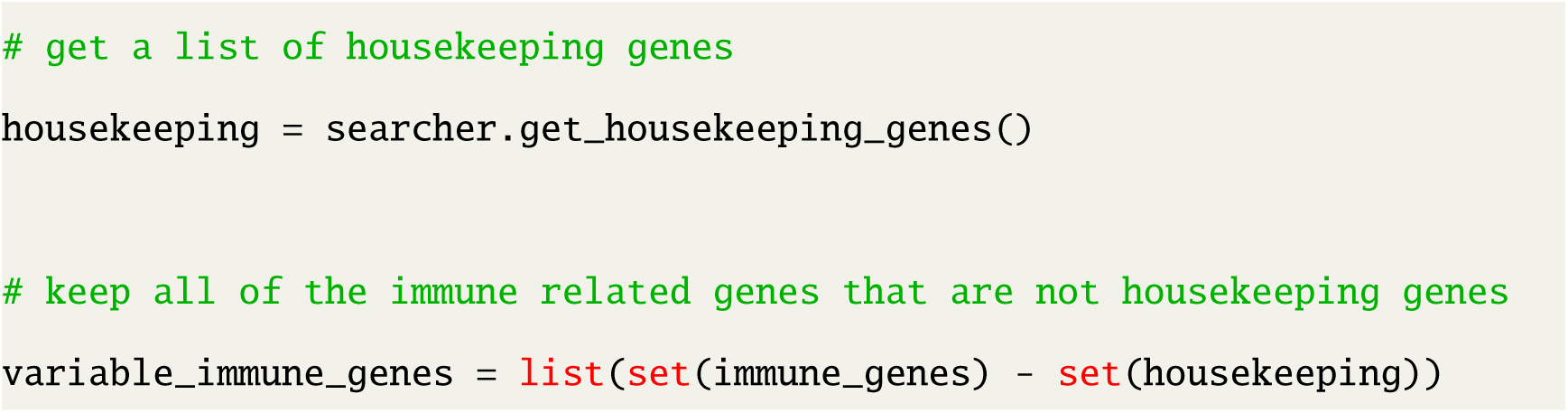

### B. Obtaining statistics about gene expression

We can use `genemunge` to access summary statistics from the GTEx project (through recount) about expression levels in healthy tissue [7–10]. The median expression value can be used to find genes that are more expressed in the small intestine than the stomach.

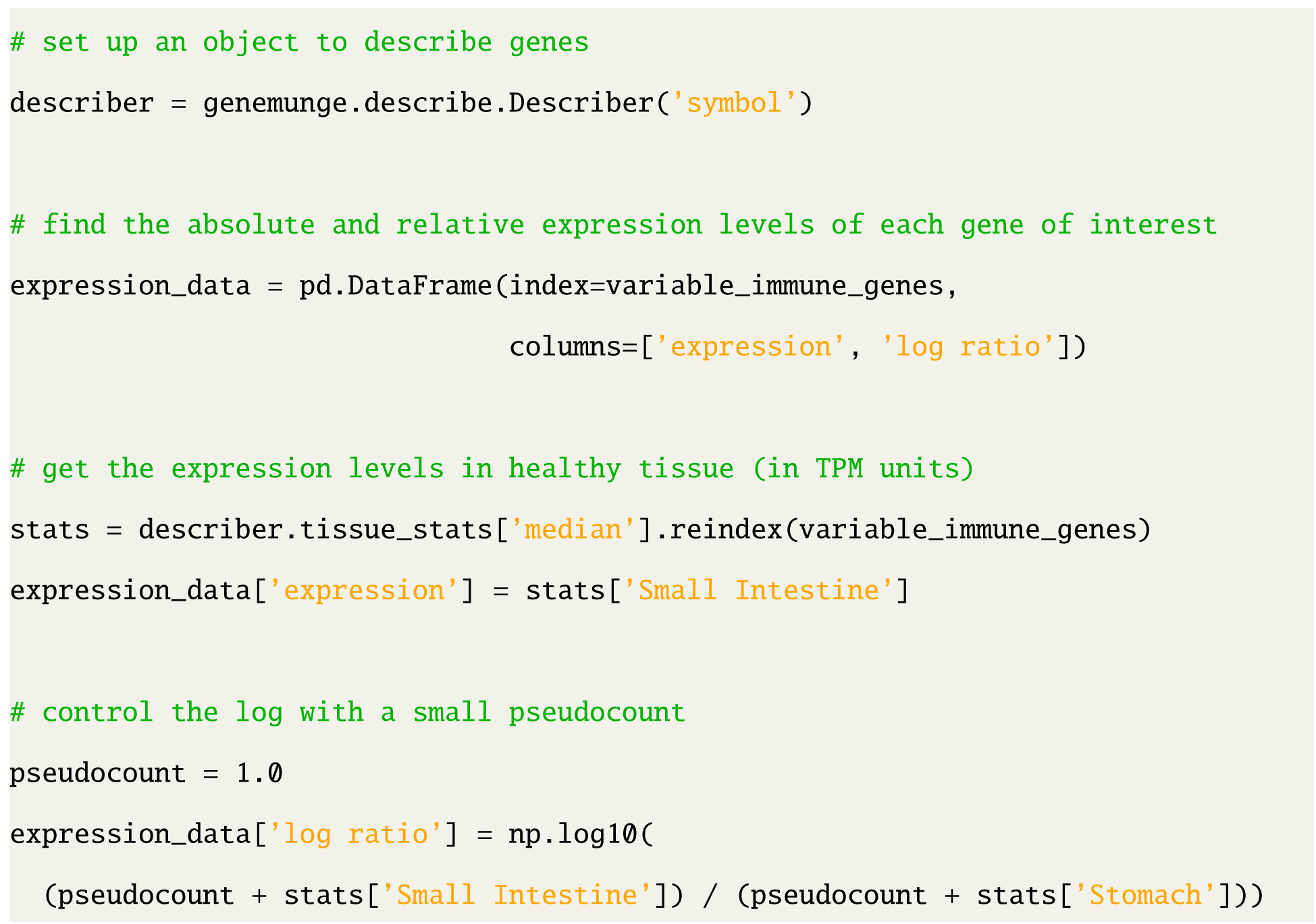

A scatter plot of the relative expression in the small intestine to the stomach against the absolute expression in the small intestine is shown in Figure 1.

**FIG. 1:**
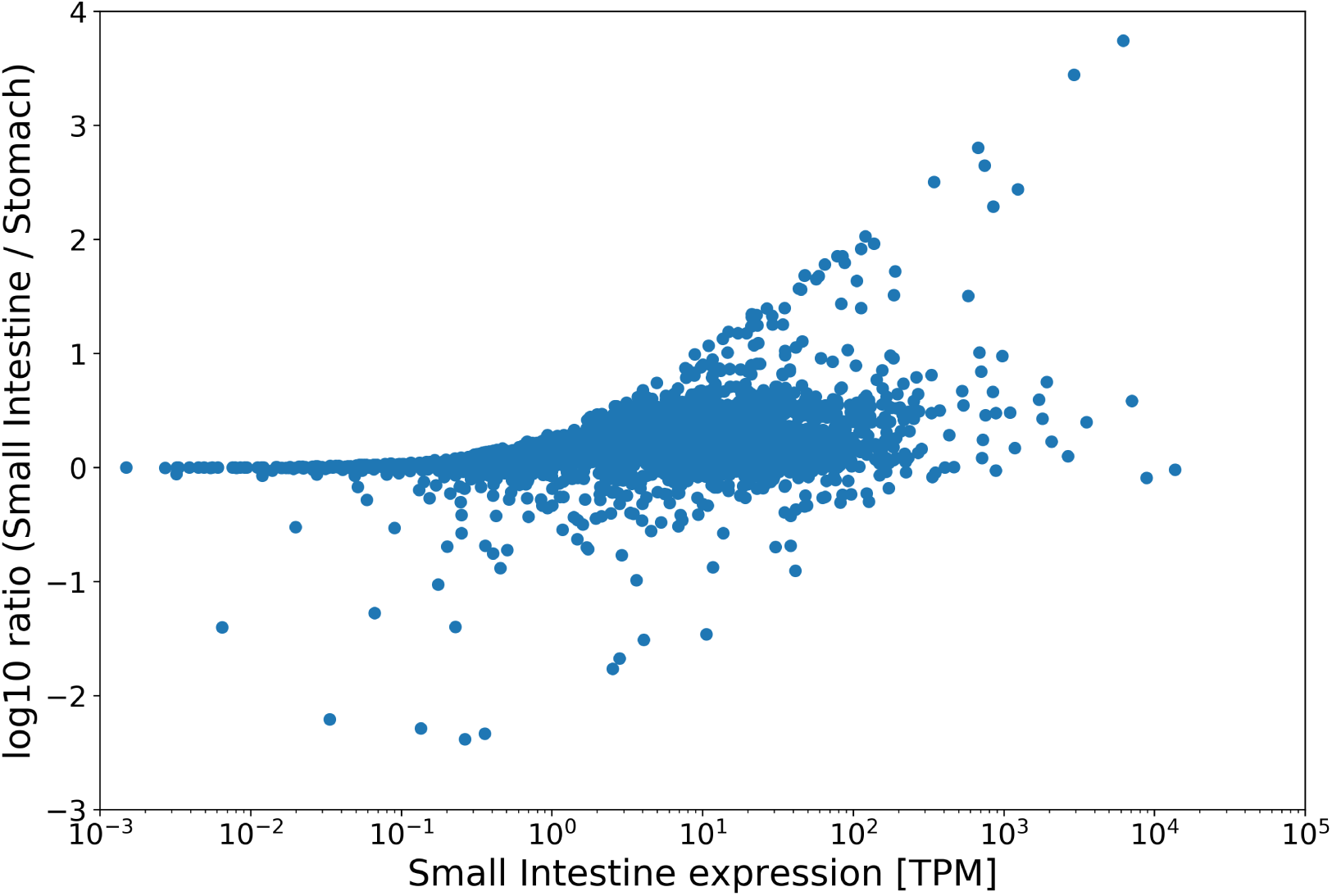
Scatter plot showing the relative expression between the small intestine and the stomach versus the absolute expression in the small intestine. Genes of interest are in the upper arm.

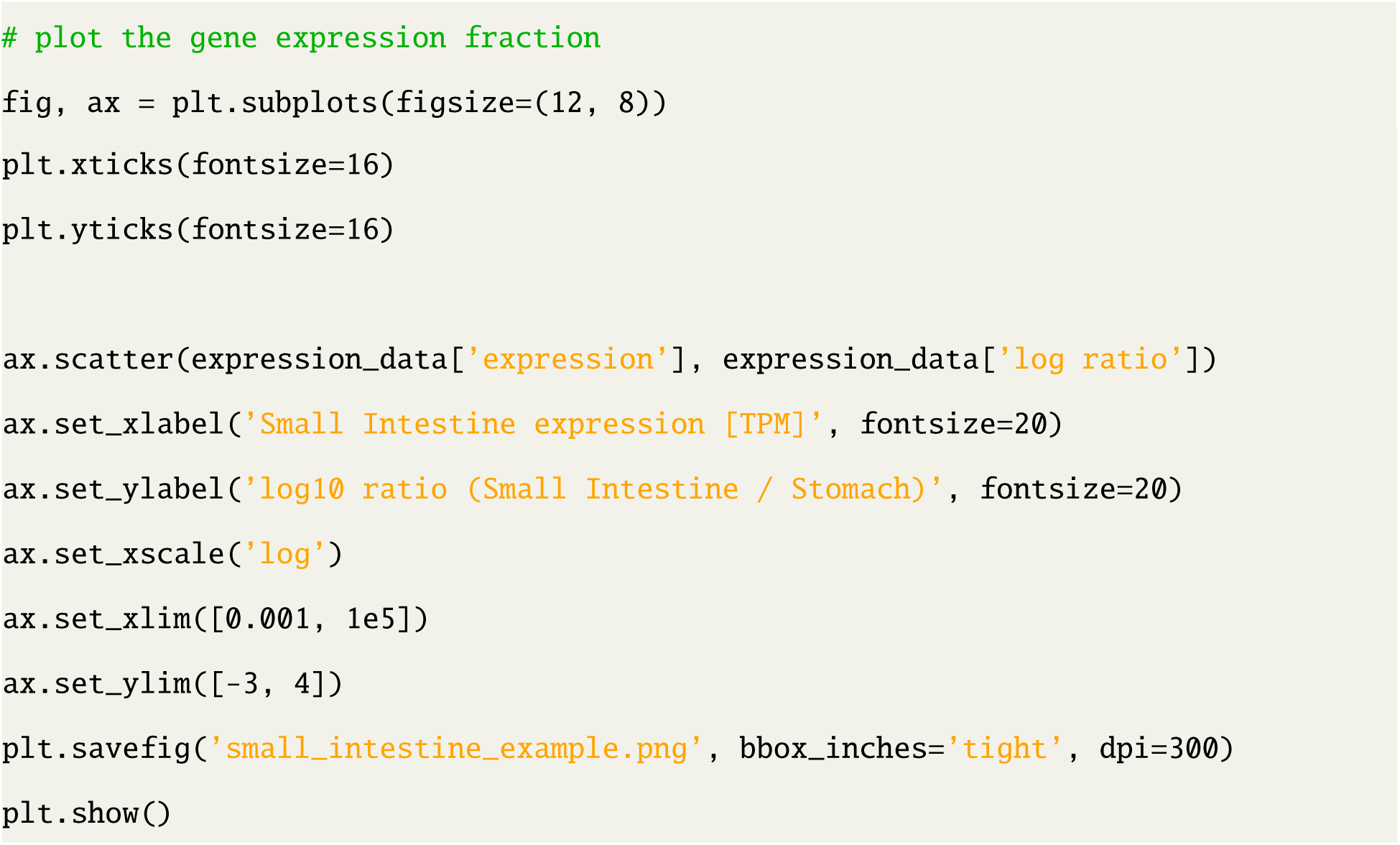

### C. Converting between gene identifier types

In `genemunge`, the base representation of genes is in terms of their Ensembl ID (without a version number). We will want to see the gene symbol in the results, so we convert the gene symbols.

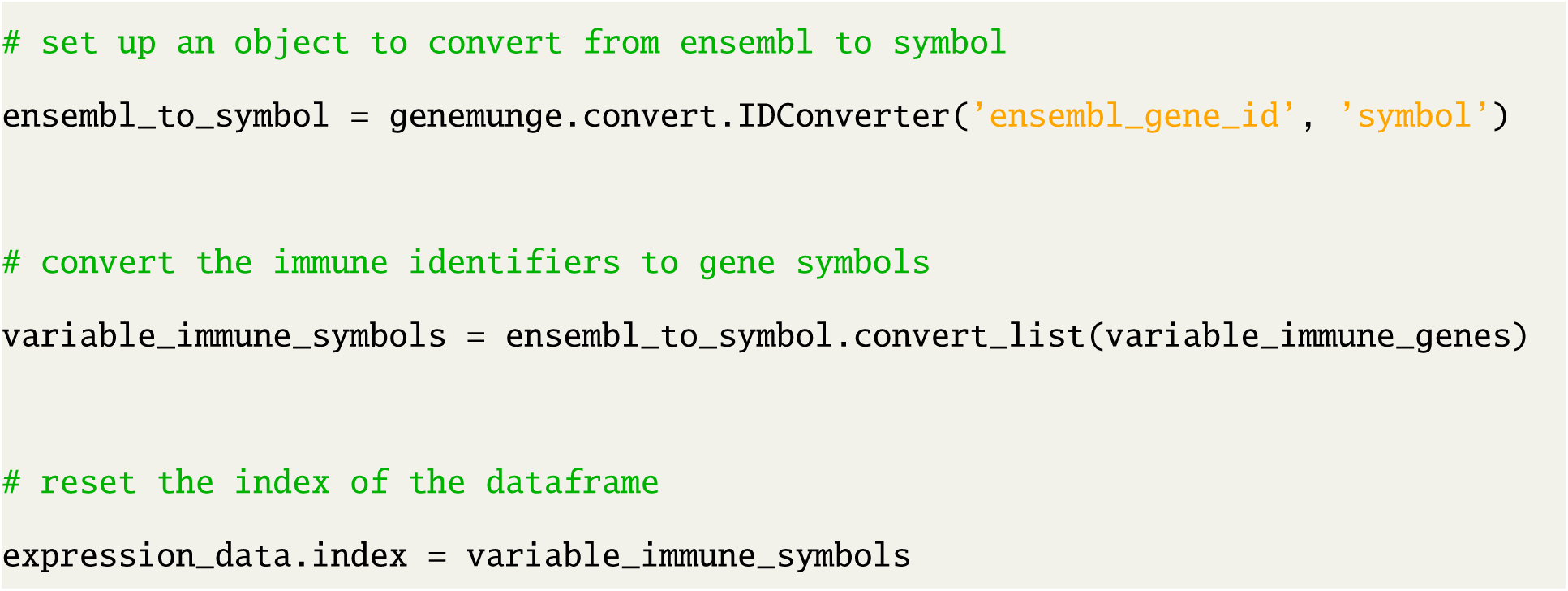

We can then select genes with high relative expression. Of course, one should be careful with this type of thing and do a differential expression analysis, but we’ll just wing it.

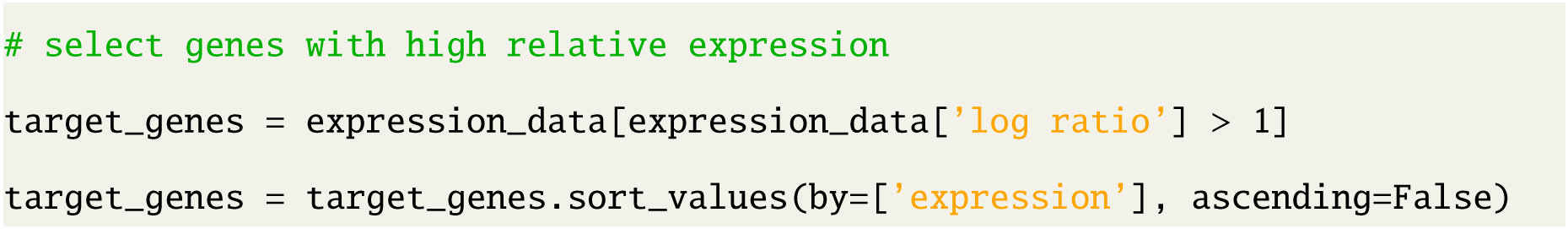

### D. Displaying information about a gene

Finally, we can look at metadata and gene expression data from GTEx on one of these genes (in this case, the most highly expressed gene).

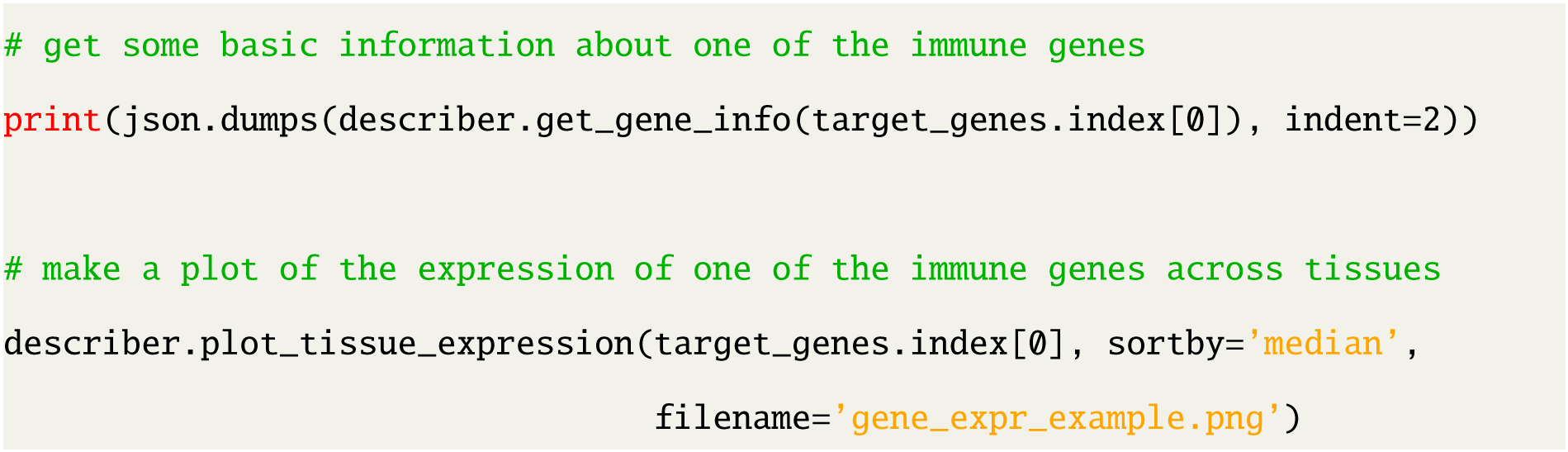

The `genemunge` output of the gene info is

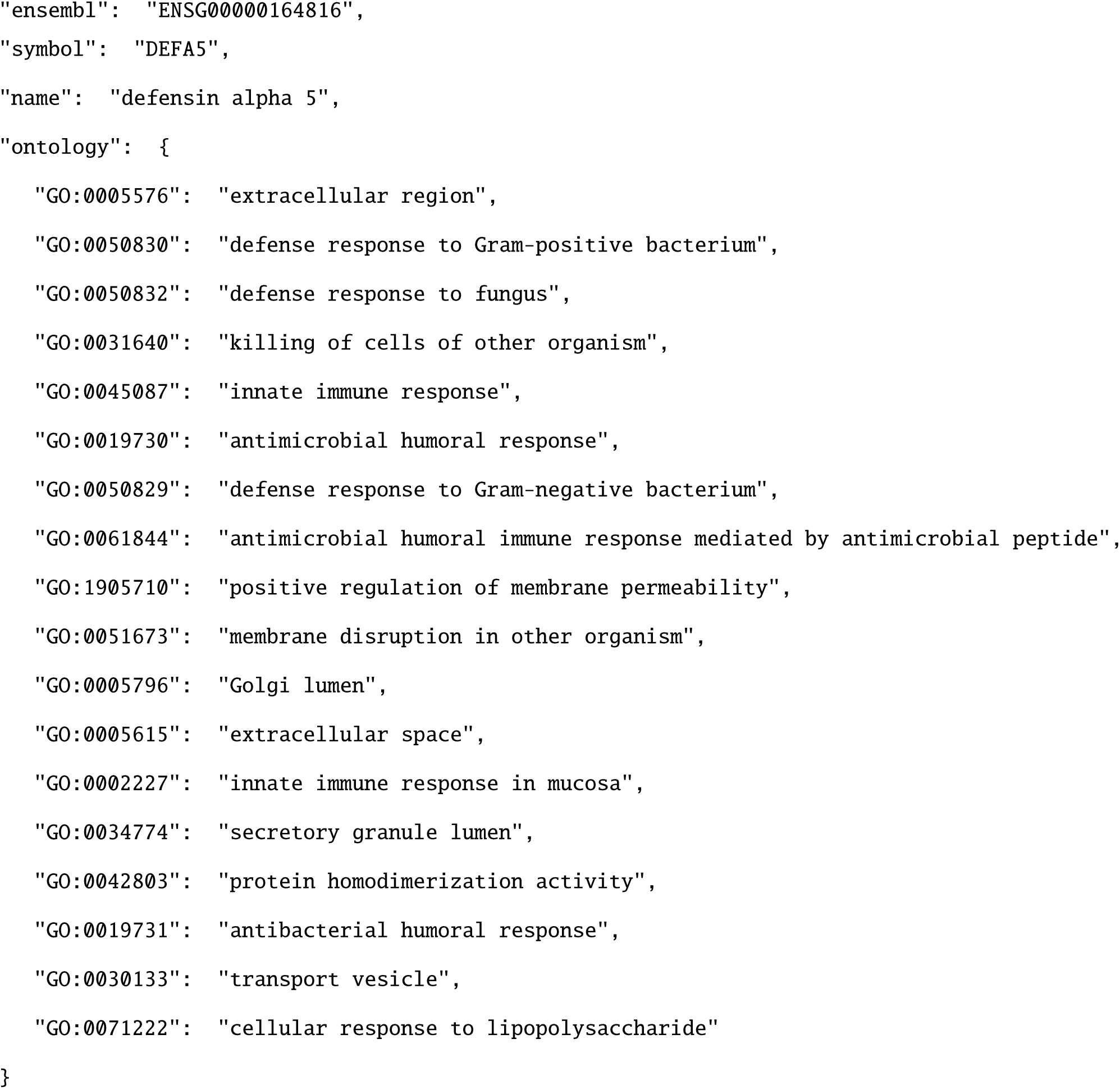

and the gene expression profile from GTEx is shown in Figure 2. As expected, DEFA5 is highly expressed in the small intestine and, in this case, nowhere else.

**FIG. 2:**
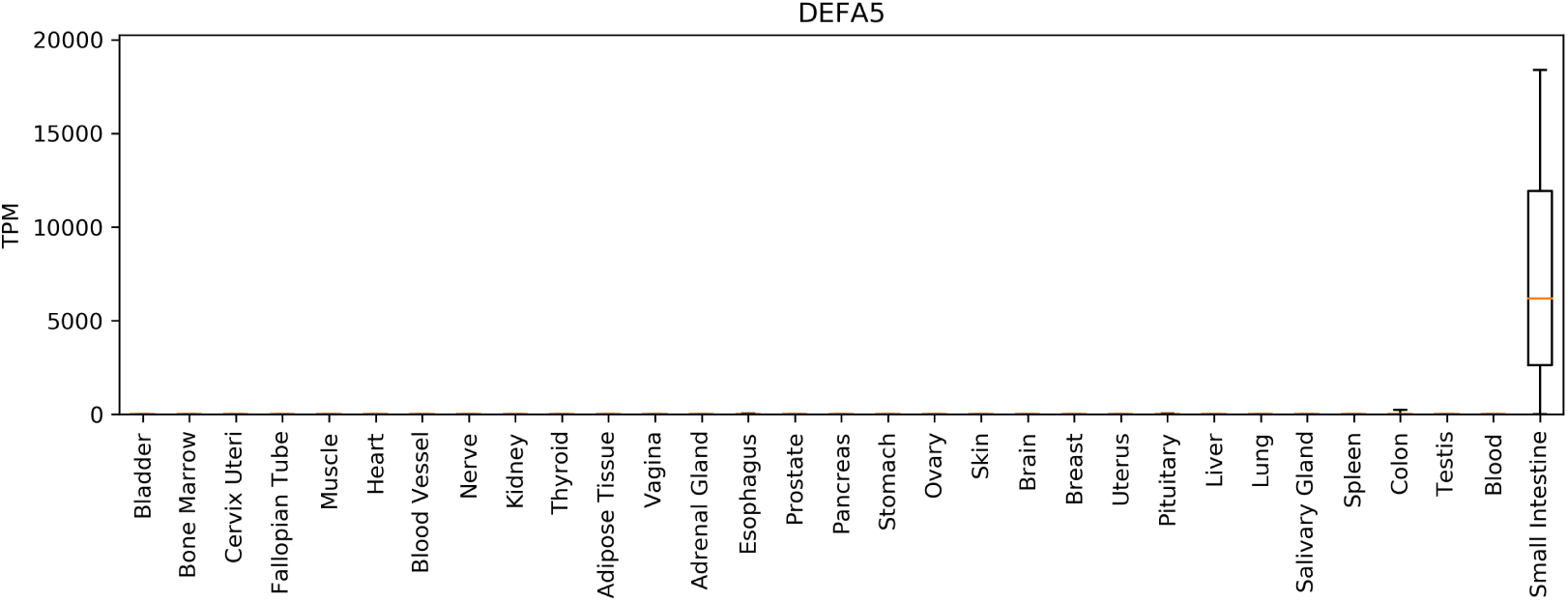
Gene expression profile in healthy tissue from GTEx for an example gene.

### E. Other stuff we aren’t covering

This example does not use all of the features of `genemunge`. For example, one could also use `genemunge` to identify transcription factors, or look up the lengths of genes and use them to convert data from counts to TPM units. You can even perform unsupervised correction of batch effects using the Remove Unwanted Variation (RUV) algorithm [12, 13]. You can read more about all of the amazing things `genemunge` can do in the documentation.

## III. SUMMARY

`genemunge` was built to make working with transcriptomics data easier. It provides a simple way to select genes of interest in an analysis and return useful metadata about them. We find that it is a useful component of a larger analysis and data processing pipeline. Our intent on open sourcing the package is to engage with the computational biology community and build it into a broadly useful tool. We welcome feedback, feature requests, and contributions on `genemunge` through `GitHub`.

## References

[1] E. A. Bruford, M. J. Lush, M. W. Wright, T. P. Sneddon, S. Povey, and E. Birney, Nucleic acids research 36, D445 (2007).

[2] M. Ashburner, C. A. Ball, J. A. Blake, D. Botstein, H. Butler, J. M. Cherry, A. P. Davis, K. Dolinski, S. S. Dwight, J. T. Eppig, et al., Nature genetics 25, 25 (2000).

[3] G. O. Consortium, Nucleic acids research 32, D258 (2004).

[4] S. Carbon, A. Ireland, C. J. Mungall, S. Shu, B. Marshall, S. Lewis, A. Hub, and W. P. W. Group, Bioinformatics 25, 288 (2008).

[5] K. Chawla, S. Tripathi, L. Thommesen, A. Lægreid, and M. Kuiper, Bioinformatics 29, 2519 (2013).

[6] E. Eisenberg and E. Y. Levanon, Trends in Genetics 29, 569 (2013).

[7] L. J. Carithers, K. Ardlie, M. Barcus, P. A. Branton, A. Britton, S. A. Buia, C. C. Compton, D. S. DeLuca, J. Peter-Demchok, E. T. Gelfand, et al., Biopreservation and biobanking 13, 311 (2015).

[8] L. J. Carithers and H. M. Moore, "The genotype-tissue expression (gtex) project," (2015).

[9] A. C. Frazee, B. Langmead, and J. T. Leek, BMC bioinformatics 12, 449 (2011).

[10] L. Collado-Torres, A. Nellore, K. Kammers, S. E. Ellis, M. A. Taub, K. D. Hansen, A. E. Jaffe, B. Langmead, and J. T. Leek, Nature biotechnology 35, 319 (2017).

[11] G. P. Wagner, K. Kin, and V. J. Lynch, Theory in biosciences 131, 281 (2012).

[12] J. A. Gagnon-Bartsch and T. P. Speed, Biostatistics 13, 539 (2012).

[13] L. Jacob, J. A. Gagnon-Bartsch, and T. P. Speed, Biostatistics 17, 16 (2015).

